## Supplementary materials for "Spiking attractor model of motor cortex explains modulation of neural and behavioral variability by prior target information"

### Supplementary Material

#### Study of metastability in an alternative network topology.

Litwin-Kumar and colleagues[1] introduced a modified architecture (termed E+I) as an extension of their E clustered network (analyzed in Fig. 2 b). This architecture incorporated a novel E to I connectivity scheme. The total population of inhibitory neurons was separated into disjunct pools where each pool of inhibitory neurons was more likely to receive input from excitatory neurons of one particular excitatory cluster. Inhibitory to excitatory connections, on the other hand, were randomly drawn from all inhibitory neurons in the total population as in the E-clustered network topology. This resulted in a dynamic reduction of the Fano Factor (FF) also in the inhibitory neuron pool (cf. Fig. 8 c in Litwin-Kumar et al.[1]).

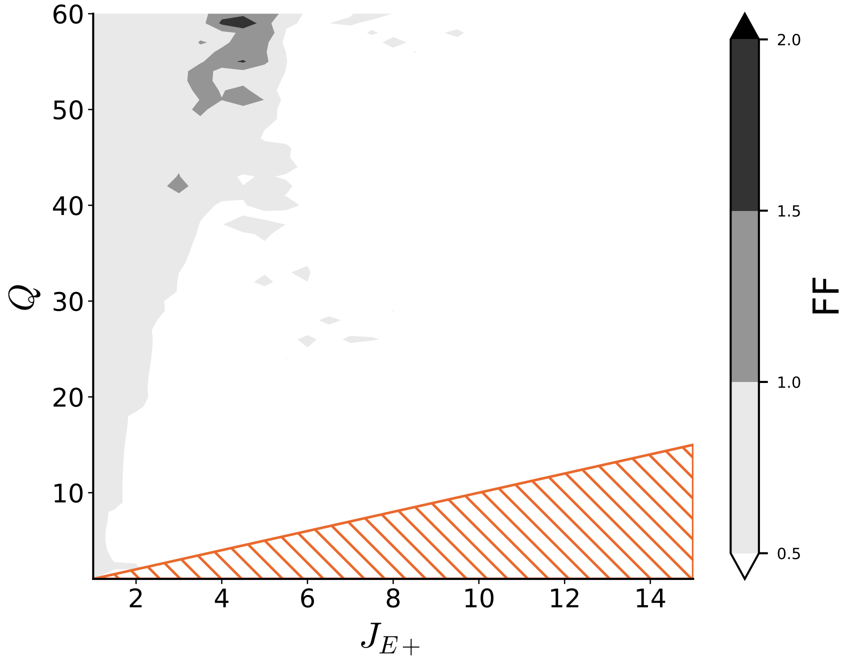


Supplementary Figure S1: **Calibration of Fano factor for extended E-clustered network[1].** Effect of cluster strength $J_{E+}$ and number of clusters $Q$ on trial-to-trial variability (FF) for the alternative E+I architecture as suggested in Litwin-Kumar et al.[1]. High FF values indicative of metastability occur only in a restricted parameter range comparable to the pure E-clustered network (cf. Fig. 2 b, main text). Shaded orange triangles indicate the zone below $J_{E+} = Q$, where the clusters are completely decoupled as in Fig. 2 b.

We re-performed our analysis of metastability (as in Fig. 2) for this topology. The results in Supplemental Fig. S1 show high FF values indicative for metastability only in a very limited parameter regime that matches the restricted parameter range obtained for the E-clustered network (Fig. 2 b). The E/I-network topology proposed in the present study is fundamentally different from the E+I network topology proposed in[1]. Our E/I model implements excitatory and inhibitory clusters that have strong mutual connections to achieve a local balance of excitatory and inhibitory input for both neuron types. This results in robust metastability across a wide range of parameters (Fig. 2d).

#### Mean-matched Fano Factor analysis.

In Fig. 5 of the main text and Fig. S3, we report the trial-by-trial variability (FF) averaged across all neurons. This analysis is based on a sliding window estimate of the empirical spike count computed for each neuron and in each single trial. For each group of trials that belong to the same experimental condition and the same final reach target we then computed the trial-averaged mean count and the FF. The parallel estimation of the spiking irregularity measured by the coefficient of inter-spike intervals (CV) predicts a certain trial-by-trial count variability across trials based on the theoretical relation of interval and count statistics in stochastic point process theory (see Methods and Discussion in the main text).

The biophysical nature of spiking neurons implies that, due to absolute and relative refractory periods that follow the generation of an action potential and in a high firing rate regime spiking can become more regular and, thus, the count variability may be reduced independent of any network property such as attractor dynamics. Therefore, and in addition to the FF in Figs. 5 and S3, we estimated the time-resolved local Coefficient of Variation ($CV_2$). We find that the population averaged $CV_2$ is constant throughout the behavioral task (Fig. 5 and Fig. S3). From this we draw two conclusions: First, the stochastic nature of the spike generating process that largely depends on the input statistics does not change throughout the task. In the E/I clustered network this is achieved by keeping the balance of excitation and inhibition constant (see Fig. 3 and Discussion). Second, the constant $CV_2$ implies that the increase in average firing rate following the preparatory stimulus (PS) does not lead to an increase in regularity and thus should not affect our measurement of the FF.

Churchland and colleagues[2] have suggested a different control to ensure that the observed reduction in FF is not affected by an increase in firing rate. Their approach is to estimate the mean-matched FF as follows. At any point in time one sample of FF is constructed from the across-trial spike count of a single neuron during identical trials (i.e. for the same task condition and the same final reach target). The trial averaged count is computed for each sample to obtain a count distribution, which is shown in Fig. S2 c and d for the unrestricted and the mean-matched analysis, respectively. In the mean-matched case, a template distribution of counts is constructed in a next step. This comprises the

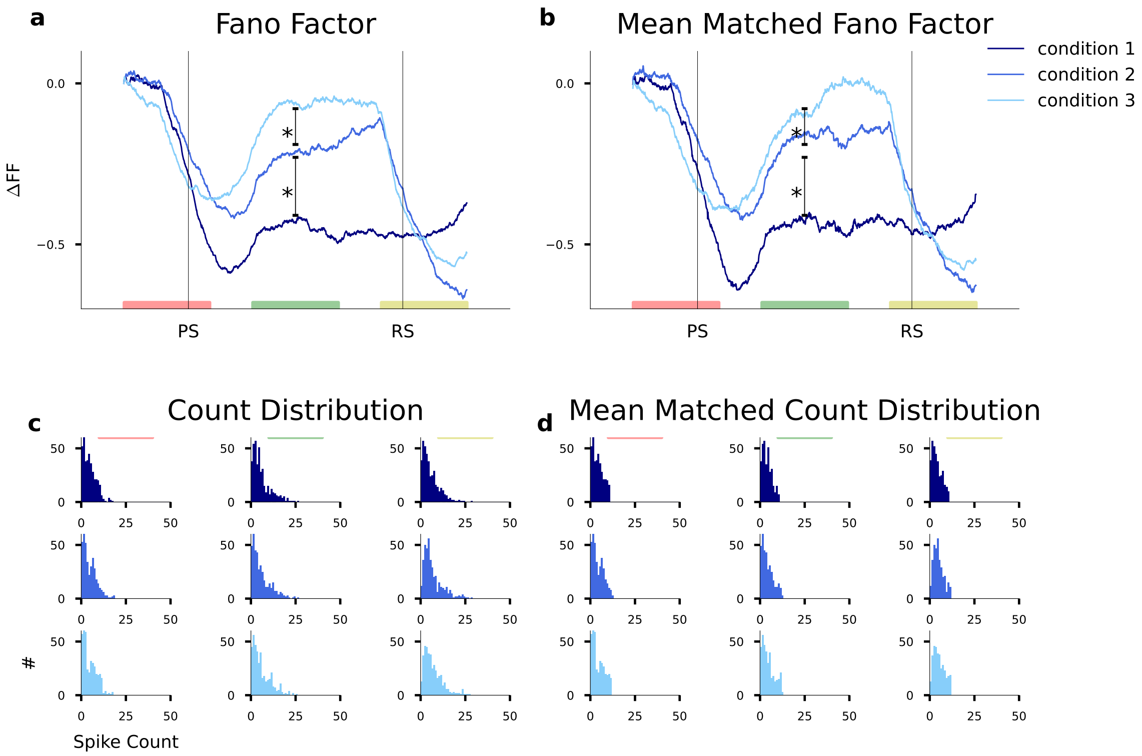


Supplementary Figure S2: **Time-resolved analysis of mean-matched FF. a)** Unrestricted analysis of FF for monkey M1 as in Fig. 5 c. Three example windows of 400 ms duration are indicated by the horizontal colored bars. **b)** Mean-matched analysis of FF for the same data as in **a** shows the same qualitative and a matching quantitative result. **c)** Empirical count distributions for the three observation windows indicated in **a** and for the three experimental conditions as indicated by the histogram color. **d)** Restricted count distributions that optimally match the template count distribution for the same three observation windows indicated in **a** and **b**.

common distribution across time constructed such that it can be matched by the sample distributions as obtained at each point in time. This distribution is considerably narrower than the distributions in the unrestricted analysis (compare panels c and d in Fig. S2) and, specifically, avoids high trial-averaged count values in phases of higher firing rates.

In Fig. S2 a and b we show the unrestricted FF analysis and the mean-matched FF analysis side by side. The mean-matched FF reproduces qualitatively and quantitatively the task-dependent reduction of FF and the significant separation for the three task conditions. In the mean-matched FF analysis, the absolute values reach even smaller values than in the unrestricted analysis, demonstrating that high firing rates are not causal for the observed FF reduction in our experimental data.

Note, that by construction the mean-matched FF analysis has its own drawbacks. Importantly, the composition of neurons (defined by the experimental condition and final target) is generally a different one at each point in time. In addition, this method is biased towards estimation of the FF from neurons that express lower rates during task performance since, as in our case, the firing rates are typically lowest during the initial spontaneous activity. This, in turn, may emphasize a potential estimation bias of the FF that can occur for very low rates[3,4]. With respect to this bias, the argumentation for the unrestricted FF analysis goes in the opposite direction. For very low rates, the bias pushes the FF towards unity as the observed process is approaching the Bernoulli process in the limit of zero rate. This bias is rapidly diminished for higher rates. Since the empirical FF in our data and model is larger than unity for spontaneous activity where average rates are lowest, any reduction of the FF in a later phase with higher average firing rates cannot be attributed to the estimation bias.

#### Analysis of experimental *in vivo* data in the second monkey (M2) and corresponding model results.

Here, we repeat our analysis of experimental data obtained in monkey M2 that performed the same task outlined in the Methods section and Fig. 5 a. Fig. S3 demonstrates our results with respect to the timed-resolved FF, firing rate, $CV_2$, and reaction times. M2 displayed a decisively different behavioral strategy. Throughout initial training and during the subsequent recording sessions M2 did not perform anticipatory movements in Condition 1 where the full target information was presented with PS at the beginning of the delay period and, thus, did not show faster reactions in Condition 1. This is signified in the almost identical reaction time distributions across all three conditions in Fig. S3 d. This is in contrast to monkey M1 that in Condition 1 showed anticipatory movements resulting in shortened reaction times (Fig. 6 b) and, in some cases, in a premature behavioral response before RS that lead to the abortion of the trial (error trial). Our interpretation is that M2 showed a reduced attentiveness to the target-specific information cued with the PS signal during the cue period and applied the same behavioral strategy in all three experimental conditions by paying attention to the RS signal before initiating a movement invariably in all three conditions (see Discussion in the main text).

The identical reaction times are paralleled in our time-resolved FF analysis of M2 that resulted in highly similar curves (Fig. S3 a). However, the FF displays a slow reduction throughout the preparatory period indicating some degree of movement preparation (see Discussion). Spiking irregularity as quantified by $CV_2$ was again constant throughout trial time quantitatively matching the spiking irregularity observed in M1.

We employed the same E/I-clustered network model as designed to reproduce neuronal responses and behavior of M1 (Figs. 5 and 6) with identical network parameters. To accommodate the distinct behaviors of monkeys M1 and M2, we adjusted only the single input parameter of stimulus amplitude (Table 2) such that during the preparatory period this was lower (reduced by 50%) as compared to the stimulus amplitude after RS. This allowed us to quantitatively capture the temporal evolution of the FF (while retaining a constant $CV_2$) observed in M2 and to produce reaction time distributions that are highly similar in all three conditions with a matching average reaction time (cf. panels d and h in Fig. S3).

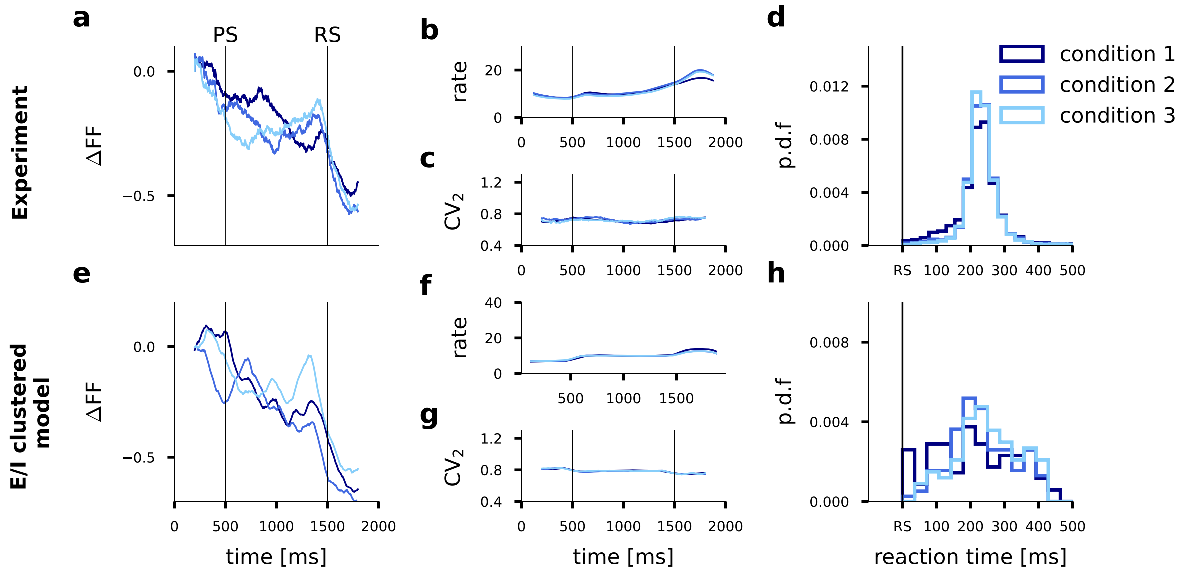


Supplementary Figure S3: **Analysis of experimental data in M2 and corresponding simulation of the functional E/I network model. a-d)** ΔFF, firing rate, $CV_2$ and reaction times calculated for 3 different conditions for monkey M2. The analysis parameters are identical to those used for monkey M1 as detailed in Fig. 5 and Fig. 6. **e-h)** Analysis of E/I network model with identical network parameters as in Fig. 5 and Fig. 6. Stimulus amplitude was adjusted as a single parameter to $I_{Stim} = 0.05\,\mathrm{pA}$ during the preparatory period and $I_{Stim} = 0.1\,\mathrm{pA}$ after PS in order to accommodate the experimental results in (a-d) for monkey M2.

### E/I network without clustering in balanced state.

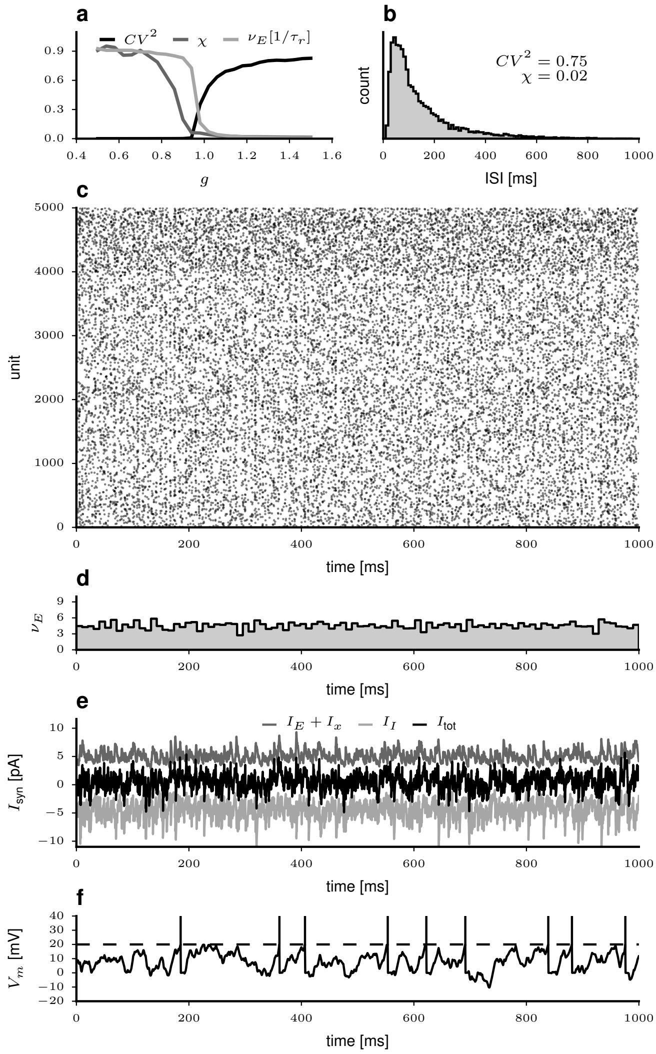


Supplementary Figure S4: **Spiking network in the balanced state.** Parameters as in Table 1 for a network without clustering ($J_{E+} = 1$). **a**) Irregularity $CV^2$, synchrony $\chi$ and normalized firing rate of excitatory neurons versus relative inhibitory synaptic strength $g$. **b**) Pooled inter-spike interval distribution for the E population. **c**) Spike raster plot during one second of spiking activity for 4.000 excitatory (Units 1 to 4.000) and 1.000 inhibitory neurons. **d**) Population rate histogram for E neurons computed in 10 ms bins. **e**) Synaptic currents of a randomly selected excitatory unit. **f**) Membrane potential for the same unit as in **e**. Vertical lines above the voltage threshold (dashed line) represent action potentials.
